# Dynamics of apomictic and sexual reproduction during primary succession on a glacier forefield in the Swiss Alps

**DOI:** 10.1101/807651

**Authors:** Christian Sailer, Jürg Stöcklin, Ueli Grossniklaus

## Abstract

Apomixis, the asexual reproduction through seeds, is thought to provide reproductive assurance when ploidy is not even and/or when population density is low. Therefore, apomicts are expected to be more abundant, and the frequency of apomictic offspring higher, at early stages of primary succession when mates are rare.

To test this hypothesis, we sampled facultative apomictic *Hieracium pilosella* L. along the successional gradient on a glacier forefield and determined their ploidy, the level of apomixis in their offspring, and the genetic diversity of the entire meta-population and within subpopulations.

We found that apomixis is more common in odd- and aneuploid cytotypes, which are more frequent at early stages of primary succession. However, apomixis was uncommon at all successional stages and sexual hexaploids were dominating throughout. Reproductive assurance was reflected in the higher fertility of all odd-ploid apomictic plants (3x, 5x) by avoiding meiosis, illustrating that apomixis provides an escape from sterility, as proposed by Darlington. Odd-ploid plants are supposedly better colonizers (Baker’s law), which is supported by their higher occurrence close to the glacier snout. Independent of succession, we found gene flow between apomicts and sexuals, which allows for the continuous creation of new apomictic and sexual genotypes.

We conclude that apomixis in *H. pilosella* does indeed provide an escape from sterility, and therefore reproductive assurance, in aneuploid cytotypes. We further propose that apomixis preserves beneficial combinations of unlinked alleles in every generation for as long as apomictic genotypes persist in the population.

## Introduction

Apomixis can be viewed as a deregulation of sexual processes, resulting in asexual reproduction through seeds^1-4^. It modifies processes central to sexual reproduction^5^: Meiosis and thus segregation is avoided (apomeiosis), and because the embryo – and sometimes also the endosperm – develops without fertilization (parthenogenesis), there is no paternal genomic contribution to the offspring. As a consequence, apomictically formed seeds are clones that are genetically identical to the mother plant.

Because apomicts do not require a mate^6^, apomixis provides reproductive assurance in obligate outcrossing plant species^6-9^. Moreover, apomixis provides an escape from sterility when ploidy is not even, such that meiosis fails^9^. In such species, apomictic genotypes are predicted to be more efficient colonizers than sexual genotypes. This view is supported by the phenomenon of geographical parthenogenesis, which describes that apomictic cytotypes are geographically more widespread than sexual cytotypes^10-14^, and the finding that invasive alien species are often apomictic^15-17^.

Although apomicts have the advantage of reproductive assurance, they are thought to accumulate deleterious mutations^18^. Without meiosis, no mechanism exists to purge deleterious mutations from the genomic pool of a population. This results in a successive reduction in fitness and, eventually, genotypes that have reached a critical threshold of deleterious mutations go extinct, a process known as Muller’s ratchet^18,19^. These considerations led Darlington to propose that apomixis is an evolutionary dead end^9^.

Nonetheless, apomixis is found in over 400 species belonging to 46 plant families^1^. This could have two major reasons: First, apomixis is a facultative, quantitative trait^1,20-22^. This means that in populations of apomictic plants also sexual individuals exist, and that apomictic individuals have residual sexuality. This enables apomictic species to purge deleterious mutations from their genomic pool, because apomicts can also, to a certain degree, reproduce sexually. Second, male sporogenesis and gametogenesis are usually unaffected in apomicts^1,23^. During male sporogenesis, apomixis loci can segregate, producing pollen that transmit genes conferring apomixis. Thus, pollen from an apomict can fertilize an apomictic (with residual sexuality) or a sexual genotype, generating new apomictic and sexual genotypes among the progeny^1,21,22,24^. As new apomictic genotypes arise from sexual reproduction, apomixis is not lost as a trait. Together, these two mechanisms provide an explanation for the high genetic variation found in apomictic populations^15,25-27^. Van Dijk and colleagues^22^ described this as the “apomixis gene’s view”, stating that apomixis persists as a trait in genotypes purged from deleterious mutations.

We chose *Hieracium pilosella* L. (mouse-ear hawkweed), a natural apomict, to study the ecological dynamics of apomixis during primary succession, i.e., the early stages of colonization of bare soil after a glacier retreat. *H. pilosella*’s endosperm development is autonomous, i.e., independent of fertilization, complying with the assumption of an advantage when possible mates are rare (conditional advantage), due to reproductive assurance^6-9^. Furthermore, apomictic and sexual genotypes can have the same ploidy level, which ranges from 3C to 8C [1C = one haploid genome]^28^. A further asset is that in *Hieracium* subgenus *pilosella* two loci, *LOSS OF APOMEIOSIS* (*LOA*) and *LOSS OF PARTHENOGENESIS* (*LOP*), have been shown to be required for apomixis^4,29,30^. The model of two independent loci explains the occurrence of four different offspring types^24^. The four offspring types are distinguished by the number of genome copies inherited from the mother and from the father, respectively. For example, offspring type 2n + n (B^III^ hybrid) means that two copies were inherited from the mother and one from the father^31,32^. *LOA* and *LOP* control two elements of apomixis, both of which are required to produce maternal clones (2n + 0, Fig. 1). If only *LOA* is present, meiosis is omitted but embryogenesis requires fertilization, leading to an increase in ploidy and paternal genomic contribution (Fig. 1). The resulting 2n + n offspring is thus generated through a mixture of apomictic and sexual processes. The same is true if only *LOP* is present, leading to offspring with reduced ploidy (n + 0, polyhaploid, Fig. 1), which is the result of meiosis and parthenogenesis, a sexual and an apomictic process, respectively. If both loci are absent, sexual reproduction occurs, leading to n + n offspring (Fig. 1). Because 2n + n, n + 0, and 2n + 0 offspring types need at least one element of apomixis for their formation, we consider them as apomictically produced offspring. In short, *H. pilosella* provides a system in which we have a good understanding of the genetic basis of apomixis and the formation of different cytotypes, allowing inferences about the processes that led to the formation of a specific individual.

**Fig 1.**
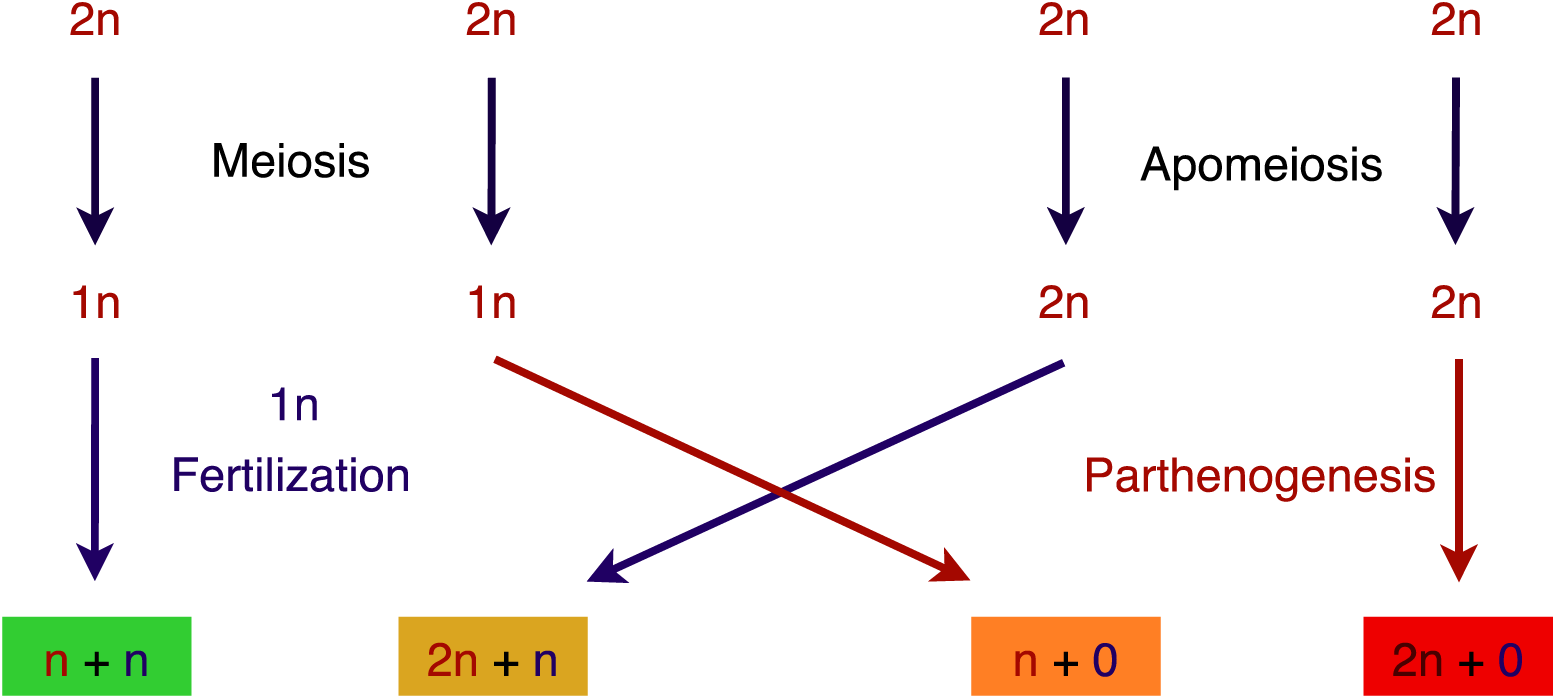
The four developmental pathways in apomictic *Hieracium* spp. The sexual developmental pathway with meiosis and fertilization generates n + n offspring (left, green). The apomictic pathway consisting of apomeiosis and parthenogenesis creates maternal clonal 2n + 0 offspring (right, red). The two loci conferring apomeiosis and parthenogenesis can segregate, resulting in mixed pathways (middle). The sexual process of meiosis combined with the apomictic process of parthenogenesis generates polyhaploid n + 0 offspring (orange). This offspring type is a new cytotype with half the maternal genomic content. The apomictic process of apomeiosis combined with the sexual process of fertilization produces B^III^ hybrid 2n + n offspring (golden). This offspring type is a new cytotype with increased genomic content, compared to the parents. Red and dark blue depict maternal and paternal contributions to the offspring, respectively.

To investigate the dynamics of apomixis and sexual reproduction, we sampled *H. pilosella* along a primary successional gradient on the Morteratsch glacier forefield in the Swiss Alps. *H. pilosella* occurs throughout the Morteratsch glacier forefield, except at the very earliest stage (Sailer C, *personal observation*). The Morteratsch forefield has a very well documented chrono-sequence of the glacial retreat^33,34^. Moreover, because of the flat topography of the forefield, we do not expect confounding influences of changes in altitude, exposition, or disturbances by avalanches and landslides on the primary successional gradient. These unique features make the Morteratsch glacier forefield a particularly well-suited model for a case study on the dynamics of apomixis along the chrono-sequence of primary succession.

We addressed the following questions concerning hypotheses of reproductive assurance of apomixis in *H. pilosella* in the glacier forefield: (1) What cytotypes of *H. pilosella* occur along the Morteratsch glacier forefield and do they differ with respect to their reproductive mode? (2) Does the relative frequency of the four possible offspring types differ between occurring cytotypes and are these frequencies influenced by the succession? In other words, does the frequency of apomicts and their level of apomixis change along the glacier forefield? (3) How have different cytotypes with different reproductive modes arisen and do they differ in their fertility?

## Results

### Apomictic cytotypes are more frequent at early stages of the successional gradient

Of the 153 plants, 142 were hexaploids. For 11 plants, we were unable to assign a ploidy level based on flow cytometry. Six of these had DNA contents between penta- and hexaploids, and five between tri- and tetraploid. Since we are unable to assign a clear ploidy level, we refer to those plants as aneuploid for simplicity. Those two cytotypes (hexa- and aneuploid) were not equally distributed along the successional gradient (2-way interaction, F_1, 23_ = 4.8, *P* = 0.039).

We found 126 plants to be sexual and 27 to be apomictic (18%), disclosing that the population on the glacier forefield consists of two reproductive types. The abundance of apomictic individuals does not change along the succession (F_1, 4_ = 2.05, *P* = 0.226, Fig. 2a; hexaploids only: F_1, 4_ = 0.057, *P* = 0.823, Fig. 2b). *Hieracium pilosella* grows in patches, often of mixed ploidy, but the majority of patches (35 of 55) we analyzed consisted solely of sexual individuals. When considering the ecological unit of a patch, we found that the frequency of apomicts within the patches decreases towards older successional stages (F_1, 53_ = 3.94, *P* = 0.052, Fig. 2c). However, this pattern is driven by 11 individuals in 4 patches. If only hexaploid individuals are considered, we did not find this trend (F_1, 52_ = 0.069, *P* = 0.794, Fig. 2d).

**Fig 2.**
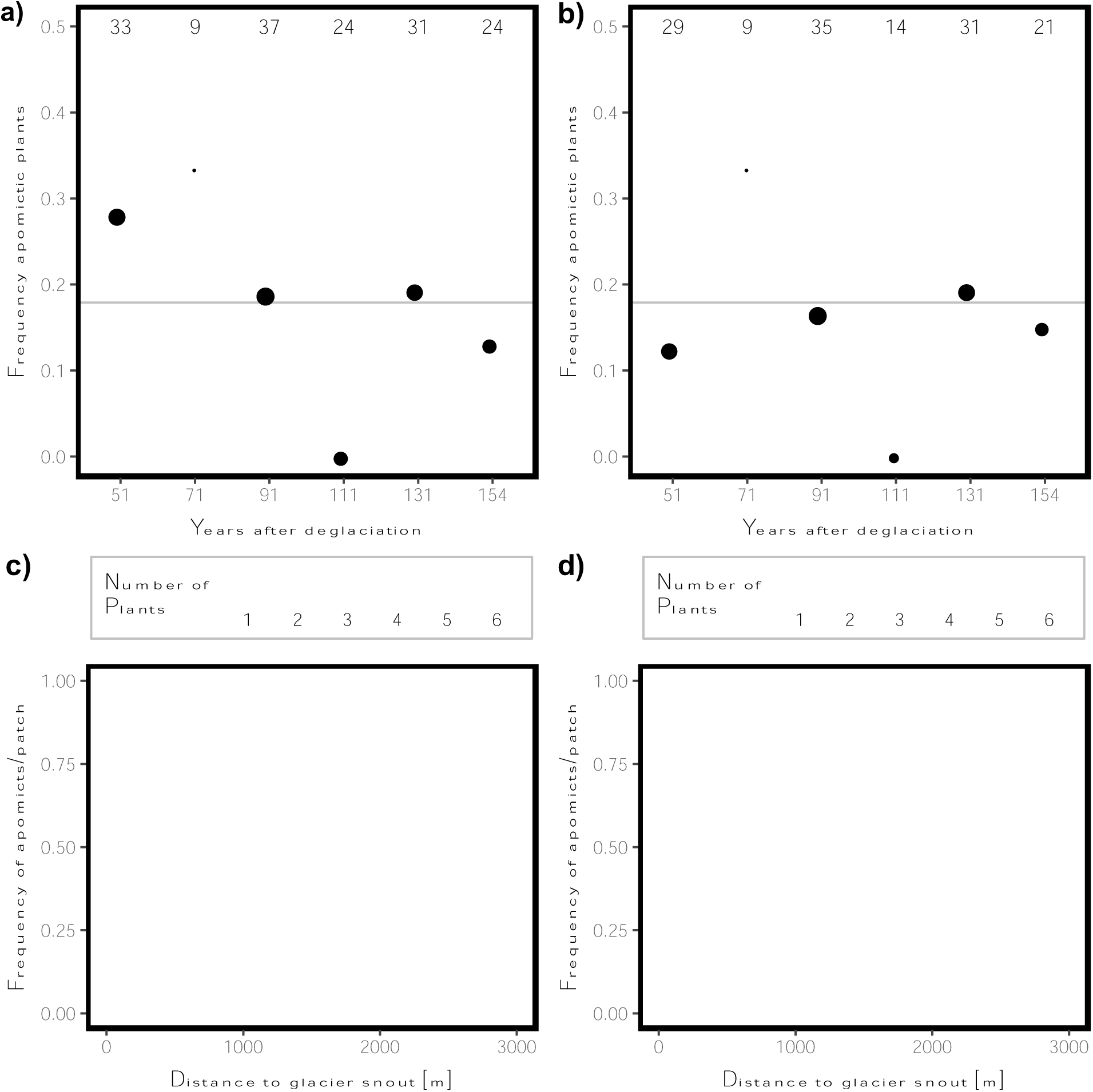
Frequency of apomicts. **a-b)** Frequency of **a)** all and **b)** hexaploid apomictic individuals in the six time windows sampled along the primary successional gradient. The frequency of apomictic plants does not differ along the successional gradient (a): F_1, 5_ = 2.05, *P* = 0.226; b): F_1, 5_ = 0.063, *P* = 0.814). The size of the dots corresponds to the number of individuals sampled. The grey horizontal line indicates the average frequency of apomicts on the forefield of the Morteratsch glacier. **c-d)** Frequency of **c)** all and **d)** hexaploid apomicts per patch along the primary successional gradient. There is a slight trend towards a lower frequency of apomicts at later stages of the succession (c): F_1, 53_ = 3.94, *P* = 0.053), which disappears if solely hexaploid individuals are considered (d): F_1, 52_ = 0.069, *P* = 0.794). 35 patches consisted entirely of sexual plants, 16 patches had apomictic and sexual individuals, and 4 patches consisted exclusively of apomictic plants. The size of the dots corresponds to the number of plants sampled per patch.

### The frequency of offspring types involving at least one element of apomixis is highest close to the glacier snout

From the total 1231 seeds analyzed, 1166 were n + n (sexual), 15 were 2n + n (B^III^ hybrid, mixed developmental pathways), and 50 were 2n + 0 (maternal clones). We did not find a single n + 0 (polyhaploid) offspring (Figure 3), indicating a bias against this specific mixed sexual (meiosis) and apomictic (parthenogenesis) developmental pathway.

**Fig 3.**
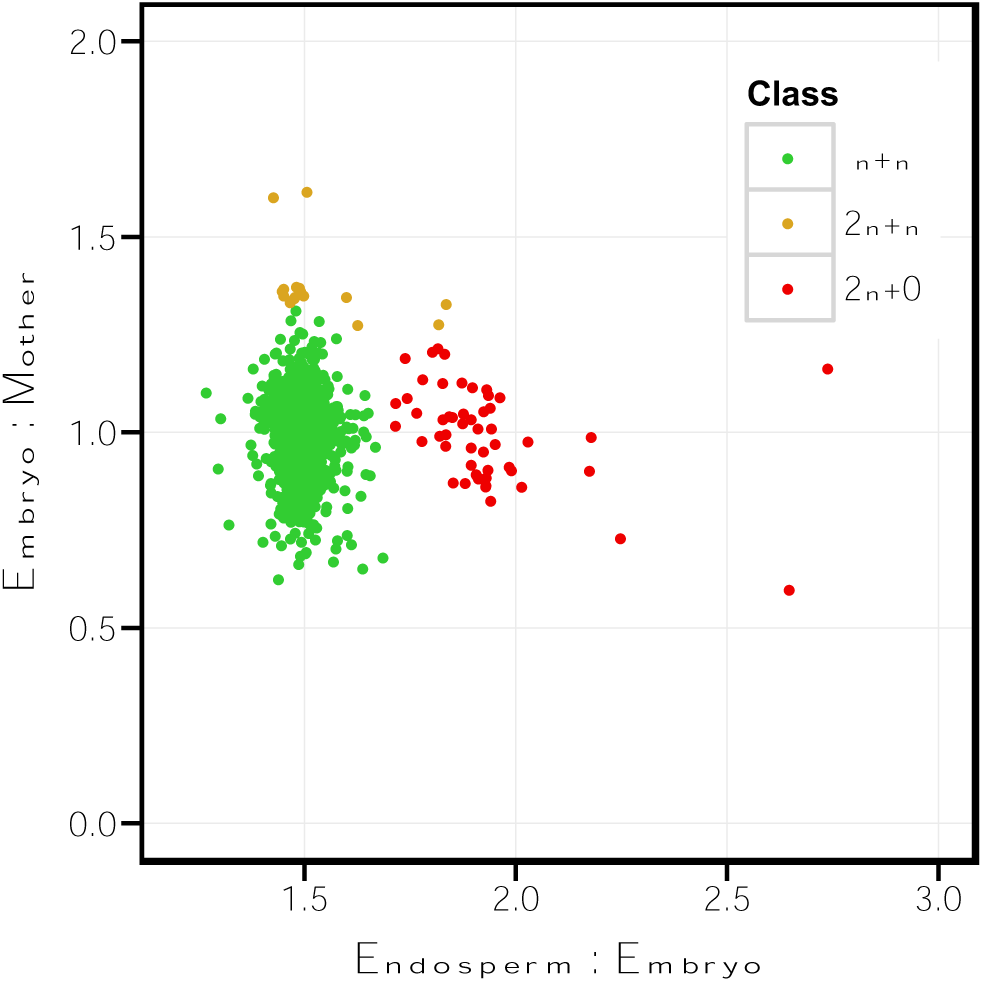
Developmental origin of seeds. Most of the 1231 seeds analyzed, coming from 153 individuals, result from the sexual pathway (n + n, green). Maternal clonal offspring, generated by the apomictic pathway (2n + 0, red), are common. Seeds produced via the mixed pathway of apomeiosis and fertilization (2n + n, golden) were rare. The fourth pathway, meiosis and parthenogenesis (n + 0), does not contribute to the seed pool we sampled. Developmental origin is determined by the ploidy ratio of embryo to mother and the ploidy ratio of endosperm to embryo. The wrong assignment rate of the linear discriminant analysis is 3.4%.

The frequency of the three occurring offspring types was mainly determined by the cytotype, i.e. ploidy of the mother plant (for 2n + 0: F_1, 1_ = 7.1, *P* = 0.015). In other words, aneuploid cytotypes had the highest frequency of apomictic offspring. Notably, apomictic hexaploid plants had a low frequency of apomictic offspring in general (14.4%), and sexual offspring prevailed in hexaploids (Fig. 4). The amount of residual sexuality varied among apomictic hexaploid mother plants (Fig. 4), illustrating the facultative nature of apomixis in *H. pilosella*.

**Fig 4.**
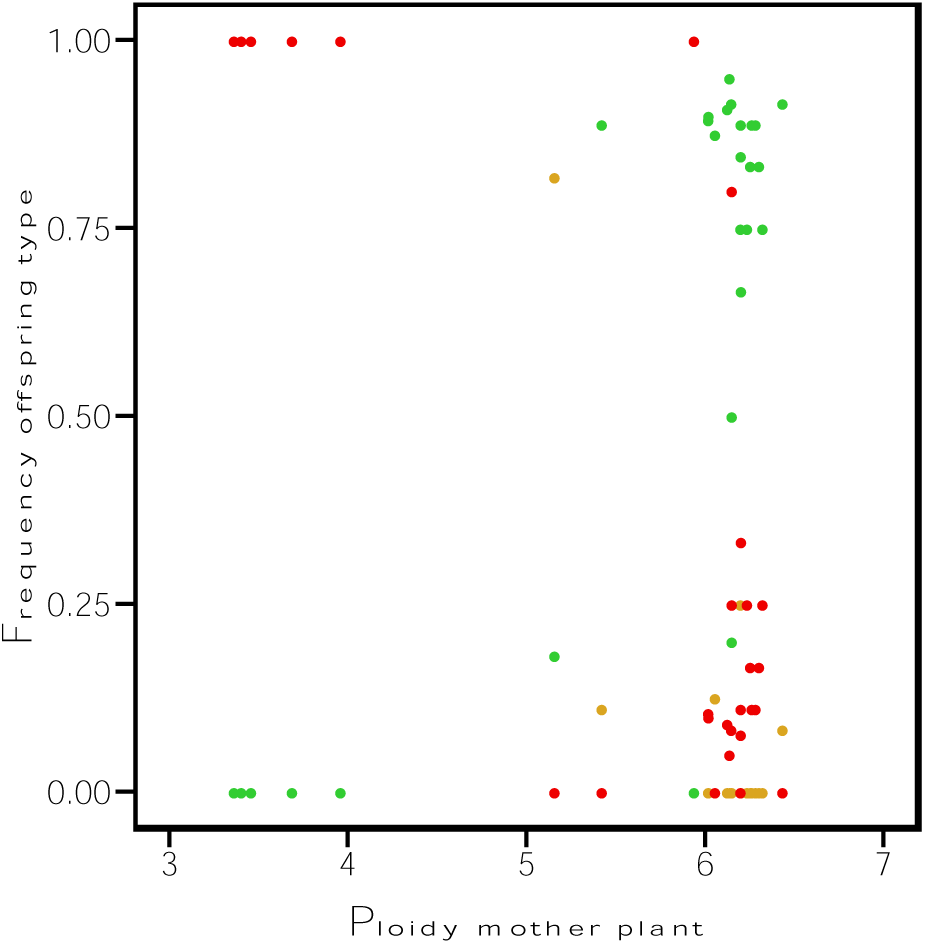
Frequency of offspring types in relation to ploidy of the apomictic mother plant. Hexaploid apomicts have a high frequency of sexual n + n offspring (green) and a low frequency of maternal clonal 2n + 0 offspring (red). Plants with an aneuploid DNA content are fully apomictic. The least frequent offspring type, 2n + n (golden), has a high frequency particularly in pentaploid plants.

Plotting the 27 apomictic plants in relation to the successional stage and the cytotype/ploidy of their mother plant revealed that the frequency of the three offspring types was unequally distributed along the succession and depended on the ploidy of the mother plant (Fig. 5). In particular, sexual (n + n) offspring from hexaploid plants were found throughout the successional gradient with a higher frequency at later stages (Fig. 5a). On the other hand, odd-ploid cytotypes had a high frequency of 2n + n and 2n + 0 offspring. Interestingly, one pentaploid plant had the highest frequency of 2n + n offspring (Fig. 5b), indicating the necessity of apomeiosis to produce seeds in odd-ploid plants. Plants with a DNA content between triploid and tetraploid produced only 2n + 0 offspring (maternal clones, Fig. 4). Remarkably, they were only found close to the glacier snout, at the earliest successional stage at which *H. pilosella* occurs (Fig. 5c). In other words, the pattern of decreasing abundance of apomictic plants in the course of succession is driven by the unequal distribution of cytotypes.

**Fig 5.**
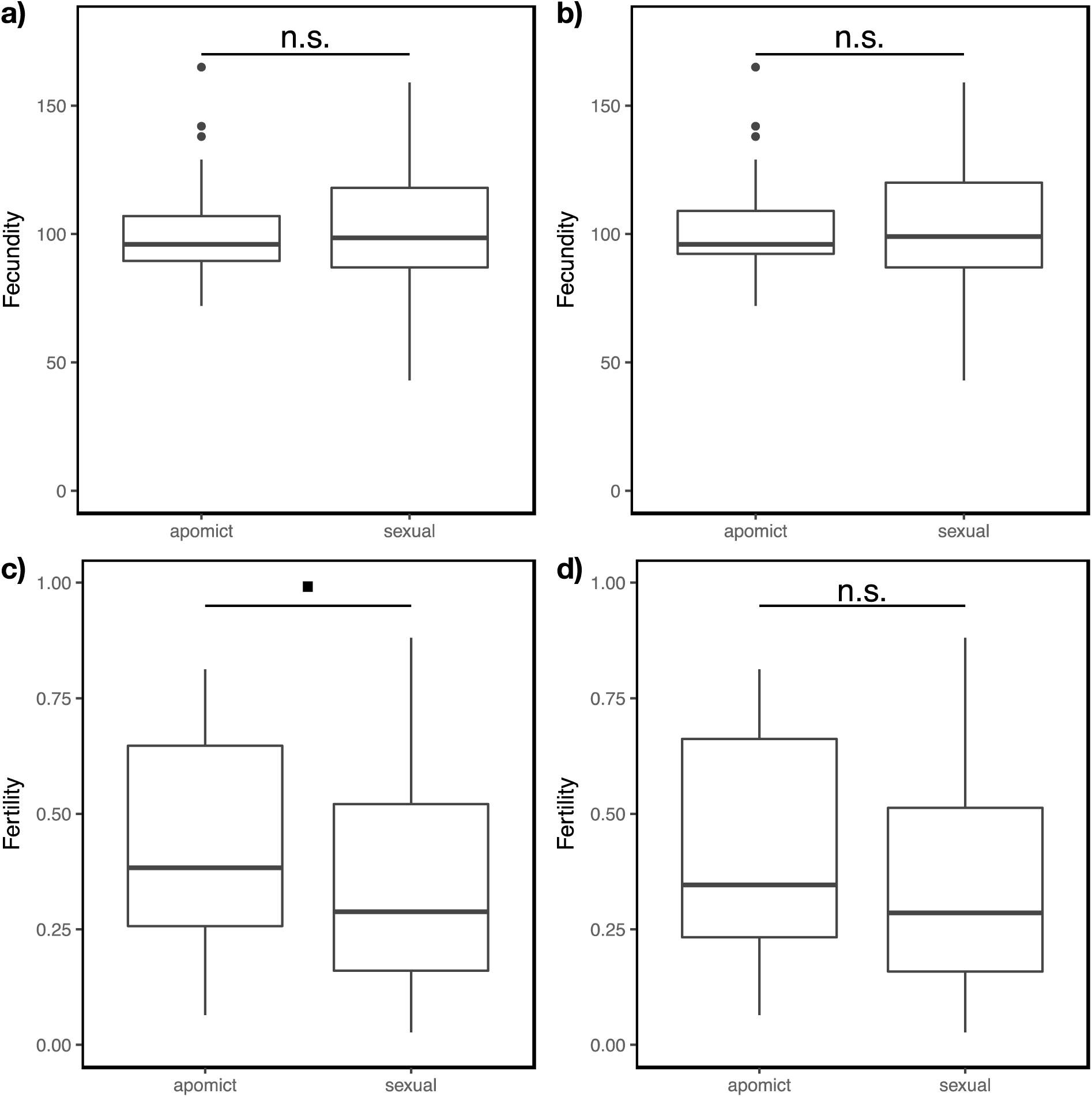
Frequency of offspring types in apomicts along the primary successional gradient. **a)** Hexaploid apomicts show a high frequency of sexual n + n offspring throughout the succession with only a few exceptions. **b)** One pentaploid apomict has a high frequency of the B^III^ hybrid 2n + n offspring. **c)** Aneuploid apomicts, which solely occur at the early stages of succession (F_1, 151_ = 12.9, *P* < 0.001), have the highest frequency of maternal clonal 2n + 0 offspring. For hexaploid plants, the frequency of 2n + 0 offspring declines along the successional gradient.

### Genetic exchange occurs frequently between apomicts and sexuals

The overall genetic diversity of *H. pilosella* on the glacier forefield was D_γ_ = 14.11. The diversity of the two subpopulations was D_apomicts_ = 11.27 and D_sexuals_ = 13.72 (Table 1). D_β_ was 1.08 (Table 1), indicating that apomictic and sexual plants cross frequently. Furthermore, we did not detect a subpopulation structure in hexaploid plants along the successional gradient (D_β_ = 1.51, Table 2), except for the apomeiosis-associated marker LOA267 (D_β_ = 2.49, Table 2).

**Table 1.**
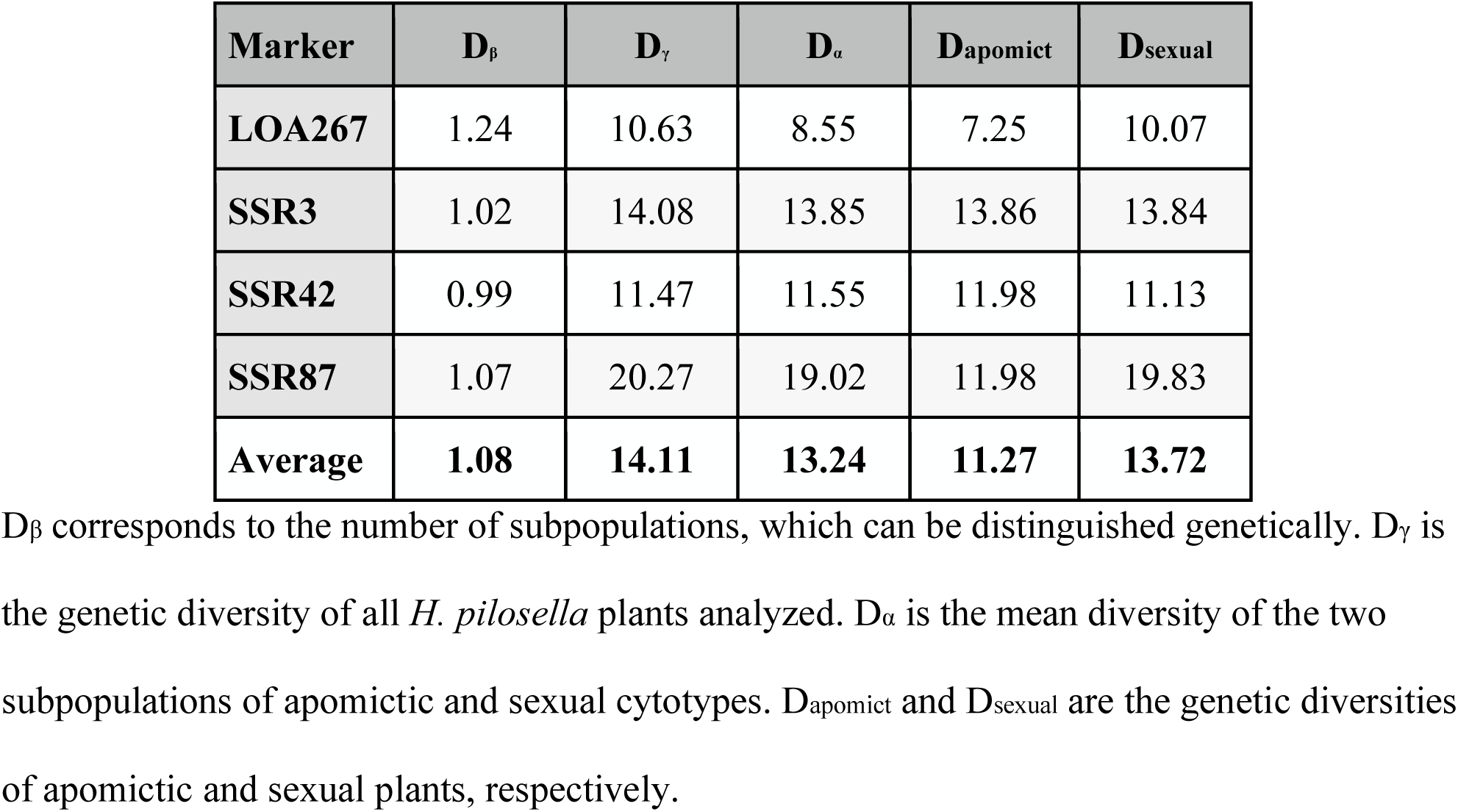
Genetic diversity of apomictic and sexual *H. pilosella* on the forefield of the Morterasch glacier, based on several molecular markers.

**Table 2.**
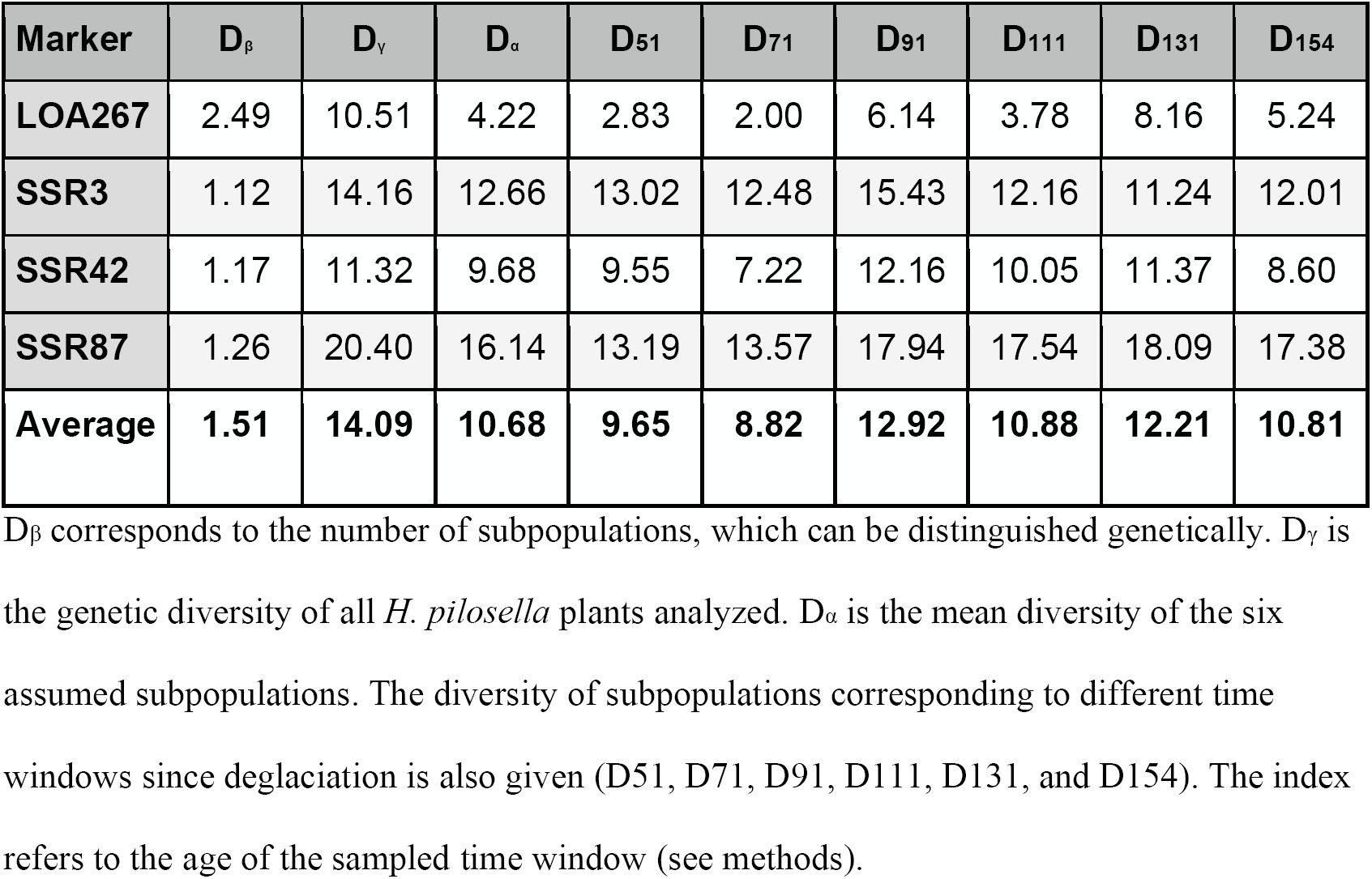
Genetic diversity of hexaploid *H. pilosella* on the forefield of the Morteratsch glacier.

Apomictic and sexual plants did not differ in their number of ovules (fecundity; F_1, 151_ = 0.09, *P* = 0.765, Fig. 6a; hexaploids only: F_1, 137_ = 0.08, *P* = 0.772, Fig. 6b), but apomictic plants had a slightly higher fertility than sexuals (F_1, 151_ = 3.6, *P* = 0.059), which was independent of succession (Fig. 6c). However, the difference in fertility is driven by the odd- and aneuploid cytotypes occurring preferentially at earlier successional stages (hexaploids only: F_1, 137_ = 2.59, *P* = 0.110, Fig. 6d).

**Fig 6.**
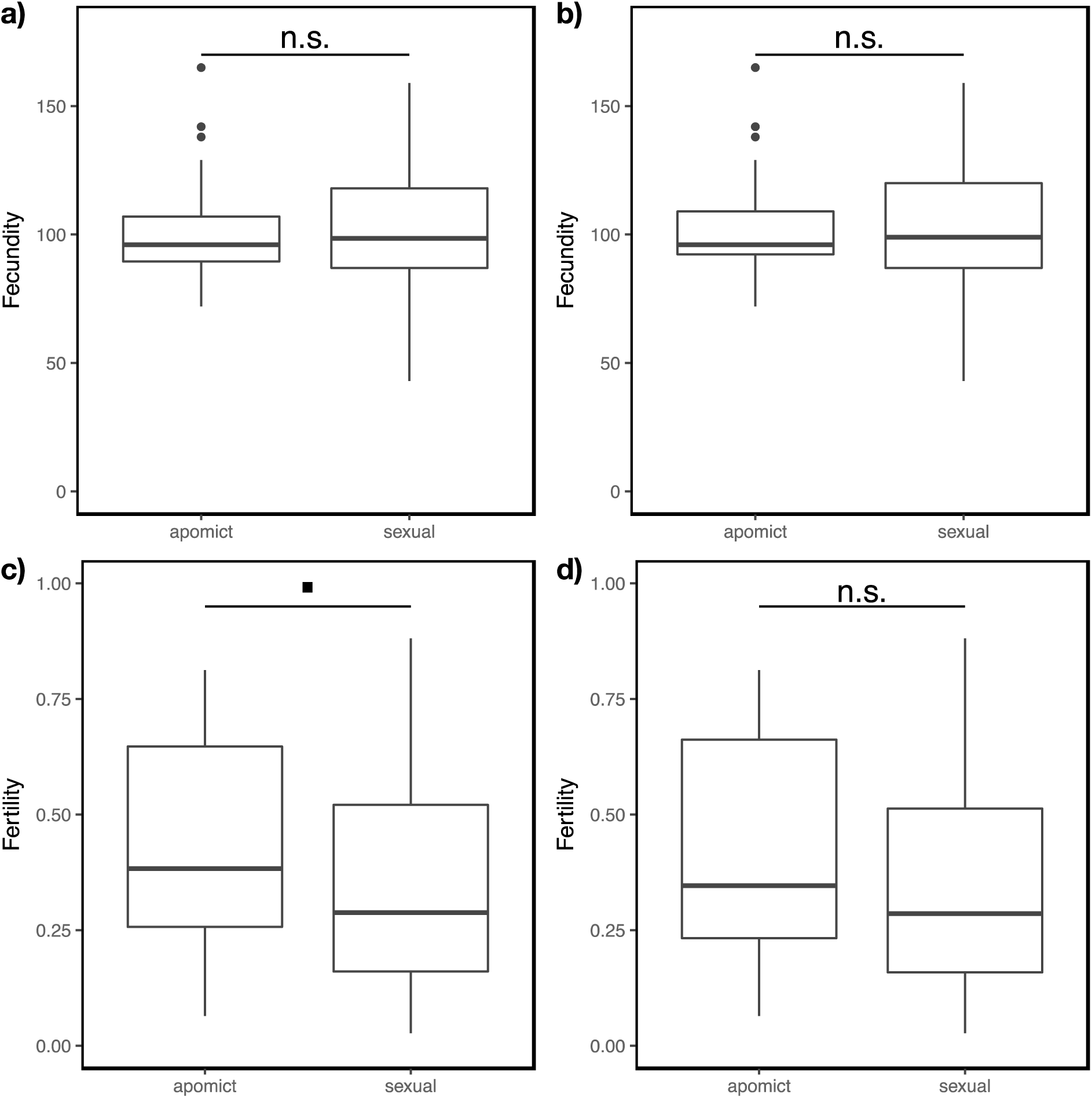
Fertility and fecundity of apomictic and sexual plants. **a-b)** Fecundity (number of ovules) of apomictic and sexual plants does not differ (a): F_1, 151_ = 0.09, *P* = 0.765; hexaploids only b): F_1, 137_ = 0.08, *P* = 0.772, b)). **c-d)** Fertility of apomictic and sexual plants. **c)** Apomicts have a marginally higher fertility (number of mature seeds/number of ovules) than sexuals, irrespective of the successional stage (F_1, 151_ = 3.6, *P* = 0.059). **d)** Fertility of apomictic and sexual hexaploid plants does not differ (F_1, 137_ = 2.59, *P* = 0.110).

## Discussion

### Different cytotypes are unequally distributed along the succession

We found aneuploid (no clear assignment of ploidy level using flow cytometry), and hexaploid cytotypes on the Morteratsch glacier forefield, and both cytotypes produced apomictic offspring (2n + n, 2n + 0). However, the majority of plants were hexaploid and produced solely sexual offspring (n + n). The identification of both sexual and apomictic offspring in the same hexaploid individuals confirms the facultative nature of apomixis in *H. pilosella.* These results are in concordance with earlier cytogeographic studies, which demonstrated the frequent occurrence of hexaploids in the Swiss Alps and described them as facultatively apomictic^1,35,36^.

Even though hexaploid plants prevailed throughout the succession, aneuploid cytotypes were unequally distributed. Cytotypes with low DNA content only occurred at early stages of succession, likely because competitive growth is dependent on ploidy, with plants being of lower plolidy being weak competitors^37^. We see hexaploids as being the more versatile cytotype in *H. pilosella* as they prevail throughout the succession and, therefore, can grow under a wide range of competitive biotic conditions.

Although only 18% of the plants were found to be apomicts, they were more frequent at early stages of succession, at which a lower density of potential mating partners is expected. For insect pollinated plants such as *H. pilosella*, mating partner density is determined by the area that is visited by a single insect. Flower visits are less frequent at early than at late stages of succession^38^, showing that mating partner density is low at early stages. While this pattern is observed if all cytotypes are analyzed together, it disappears if only hexaploid individuals are analyzed. In other words, the higher abundance of apomicts at early stages is driven by the higher abundance of aneuploid cytotypes.

Therefore, our finding of a higher frequency of apomicts at early stages of succession does not comply with Tomlinson’s model^6^, which states that selfing is prevailing when mating partner densities are low, and its interpretation that apomicts have a conditional advantage when mating partner density is low.

### The frequency of apomictic offspring is mainly influenced by ploidy level

We found a continuous variation of the frequency of apomictic offspring (residual sexuality) in hexaploid individuals, confirming that apomixis can be viewed as a facultative, quantitative trait even in predominantly sexual cytotypes. The high frequency of sexual offspring is in concordance with the sexual developmental pathway being the default in *Hieracium* spp.^4^, supporting the view of apomixis as an acquired gain-of-function trait.

Furthermore, we found that different cytotypes produced different ratios of the four possible offspring types. Plants with the lowest DNA content (< tetraploid) solely produced 2n + 0 offspring and plants with a DNA content between penta- and hexaploid 2n + n offspring, respectively. The ‘2n’ indicates the apomeiotic origin of these offspring (Fig. 1), which complies with plants of odd ploidy or aneuploidy being able to produce seeds only if meiosis is avoided^1,15,23,26,30,39^. The avoidance of meiosis provides an escape from sterility, as Darlington stated^9^, a view that is supported by our results because the vast majority of offspring were of apomeiotic origin in these aneuploid plants. They themselves, however, are likely the product of the n + 0 offspring type.

In general, we found low levels of apomixis on the glacier forefield (18%), indicating little advantage for apomicts during primary succession. The observed low level of apomixis is in concordance with earlier findings on apomictic species in the nival zone of the Alps^40^. However, *H. pilosella* plants are capable of reproducing via vegetative stolons. Like apomixis, this enables clonal reproduction, both for apomictic and sexual genotypes, although offspring number and dispersal distance are limited. We speculate that the general advantage of apomicts in *H. pilosella* is confounded by clonal reproduction via aboveground stolons of both sexuals and apomicts.

Another deduction from Tomlinson’s model^6^ is that apomicts at early stages of succession should have a high frequency of apomictic offspring. Indeed, we found more apomictic offspring (2n + 0, 2n + n) near the glacier snout. However, only aneuploid cytotypes had a high frequency of apomictic offspring, and different cytotypes are not equally distributed along the primary succession as described above. Our results suggest that the decrease in frequency of apomictic offspring as succession proceeds is primarily driven by the decrease in the frequency of cytotypes that produce high levels of apomictic offspring. Together with the high variability of the frequency of offspring types in hexaploid plants, we conclude that the level of apomixis is mainly determined by the genetic factors ploidy and reproductive type.

### Apomixis provides a source of beneficial allele combination in the formation of new cytotypes

Figure 1 illustrates that combining sexual and apomictic developmental pathways results in a change of ploidy and can explain the generation and occurrence of different cytotypes. However, progeny from such mixed developmental pathways are expected to be rare^24^. Indeed, we found not a single n + 0 offspring among more than 1200 seeds screened. Moreover, we would expect a strong selection pressure against decreasing ploidy, as two genome copies are the minimum for successful meiosis and (partial) hemizygosity can uncover deleterious alleles. As a consequence, genotypes that mainly produce n + 0 offspring will be selected against, which is likely the reason why we did not find a single n + 0 offspring in the field. Such negative selection is not expected for 2n + n offspring, which results in a ploidy increase.

Despite the rarity of n + 0 offspring, we found two patches of plants with a DNA content that was close to triploidy. These can only arise from n + 0 offspring, as most of the plants in the Morteratsch population are hexaploid. We consider each of these two patches to be a rare developmental and demographic event, as the n + 0 offspring bearing seed had to germinate and survive until maturity and reproduction. Taken together, we interpret this observation as support for Baker’s law, which states that a single individual which is capable of self-reproduction, is sufficient to found a new population^7^.

The high frequency of sexual offspring (n + n) in hexaploid apomicts throughout the succession hints towards frequent genetic exchange. Based on genetic diversity data, we found that apomicts and sexuals behaved like a single population (D_β_ close to 1), suggesting random mating between individuals with different modes of reproduction. This enables the generation of new apomictic and sexual cytotypes and supports the ‘apomixis gene’s view’^22^ while contradicting Darlington’s ‘dead end of evolution’ hypothesis^9^, at least for as long as facultative sexuality exists in this species.

Apomixis fixes genotypes and if an apomictic genotype is successful in the sense of growth and reproduction, beneficial allele combinations are frozen. These allele combinations are provided in every generation to the population’s genomic pool via gene flow between apomicts and sexuals. In contrast to sexual reproduction, the beneficial allele combinations of successful apomictic genotypes, which can cover large parts of the genome, are not broken down by recombination. We propose that apomixis preserves successful genotypes, which can repeatedly serve as a source of beneficial combinations of unlinked alleles in every generation for as long as the genotype persists in the population.

We estimated genetic diversity along the succession by assuming six subpopulations corresponding to the six time windows sampled. As we found no strong differentiation among the subpopulations from different time windows on the glacier forefield, based on neutral SSR markers (D_β_ close to 1), we conclude that the genetic exchange (gene flow) along the successional gradient is high. In contrast, the presumably non-neutral LOA267 marker, which is associated with apomeiosis, showed low diversity at early stages of succession and the D_β_ value suggests 2-3 subpopulations. Taken together, this suggests a cline along the successional gradient, pointing towards less genetic diversity near the apomeiosis locus at early stages of succession, in which apomicts are more frequent. The lower diversity at this non-neutral marker is a signature of selection for apomeiosis at these early stages, further supporting a selective advantage of apomixis at early stages of succession. Because recombination around the locus controlling apomeiosis is suppressed in most apomicts^5^, LOA267 likely reflects the segregation of a larger genomic region. However, given the current genotyping methods for *H. pilosella* and the yet unidentified genes conferring apomixis, interpretations based on the LOA267 marker alone remain speculative.

## Conclusions

We found a higher frequency of apomictic *H. pilosella* at early stages of primary succession on the Morteratsch glacier forefield. This higher frequency is due to the higher abundance of aneuploid cytotypes that do have the highest level of apomixis in this meta-population. Apomixis does provide an escape from sterility and reproductive assurance for such cytotypes, which themselves are likely to be the product of an apomictic developmental pathway. We conclude that the primary conditional advantage for apomicts is not necessarily the low density of potential mates but rather the escape from sterility for odd- and aneuploid cytotypes.

## Materials and Methods

### Model species and sampling

*Hieracium pilosella* L. is a self-incompatible, perennial, monocarpic, stoloniferous, herbaceous species. *H. pilosella* usually grows in patches of individual plants (rosettes). When rosettes reach a threshold size, they reproduce vegetatively via aboveground stolons and through seeds by producing a single flower head on a stem^41^. *H. pilosella* occurs in ploidy levels from 3C to 8C, with 5C cytotypes found at the margins of its geographical occurrence and 6C cytotypes being found throughout Europe, predominantly in the Alps [84% in Switzerland]^42^. Although *H. pilosella* is an obligate outcrosser, self-pollen germinates if non-self-pollen is also present on the stigma [mentor effect]^35^.

In preparation for sampling, the whole Morteratsch glacier forefield of ca. 1.5 km^2^ was searched for occurrence of *H. pilosella* and the positions of 912 patches were marked with GPS (GPSmap 60CS, Garmin, Garching, Germany) to an accuracy of 5 m. The positions were transferred to the topographical Swiss map (Topo Schweiz V1, Garmin, Garching, Germany) using the MapSource software (Garmin, Garching, Germany). The map with the marked positions was printed and the data of the chrono-sequence of deglaciation, dating back to 1857 when the Morteratsch glacier had its maximal extent^43^, was constructed based on a published map^34^. The glacier forefield was then sub-divided into six twenty-year time windows (51, 71, 91, 111, 131, and 154 years after deglaciation). Patches of *H. pilosella* lying on the isochronal lines, i.e., lines connecting the glacial front at certain years, as published by Burga and colleagues^34^ were dismissed. Per time window, ten patches of *H. pilosella* on each side of the river were randomly selected for sampling. From each of these patches, we aimed at collecting six reproducing plants. Sometimes, we could not find six flowering plants per patch and sampled all occurring reproducing plants in the patch instead. Furthermore, some seed samples were lost. For analysis, we only used plants from which we could sample DNA from leaves and seeds. Leaves for DNA analysis and seeds from flower heads could be collected from 234 mother plants, coming from 74 patches. In July 2011, the two youngest leaves of each individual were sampled for ploidy determination and DNA extraction. One leaf was shock-frozen in a vapor-shipper (SC 4/2 V, MVE Biomedical, Georgia, USA). The tip of the second leaf was placed in a 1.2 mL cluster tube (Thermo Scientific, Wohlen, Switzerland) containing 50 µL of mQ water (conductivity > 18 MΩ^-1^) and one 3 mm stainless steel bead (Schieritz & Hauenstein AG, Zwingen, Switzerland), and stored in a cooling bag. Closed capitula from the same plants were bagged using individually marked tea filters for seed collection. In August 2011, the individually marked tea filters containing the seeds were collected and placed in plastic containers containing silica gel to ensure fast drying of the seed material. Seeds were stored at 4°C, 30% humidity until used.

### DNA extraction and genetic diversity estimation

DNA was extracted from the sampled leaves of mother plants using the DNeasy Plant Mini kit (Qiagen, Hombrechtikon, Switzerland), following the manufacturer’s instructions. Samples were eluted in 2 x 50 µL AE buffer.

We used markers for *LOA* and *LOP*^29,44^, as well as SSR markers for *H. pilosella*^45^, to estimate overall genetic diversity of the entire meta-population on the glacier forefield. All primer pairs were tested and optimized for our samples. SSR markers were resolved on the high-resolution cartridge of the Qiaxcel system (Qiagen, Hombrechtikon, Switzerland). Three SSR markers and one *LOA* marker were highly polymorphic and could be used for genetic diversity estimation using Shannon’s entropy^46^. We calculated H_γ_ as the overall genetic diversity of the Morteratsch population. We considered apomicts and sexuals or the samples from the six different time windows as subpopulations. Using apomicts and sexuals as subpopulations, we could test for gene flow between the individuals of different modes of reproduction. Using the individuals from the six time windows as subpopulations, we could test for changes of genetic diversity along the primary succession. H_α_ was computed as the mean diversity of the subpopulations. H_β_ was computed as H_γ_ minus H_α_. H_β_ is interpreted as the number of subpopulations present in the population, based on genetic diversity. Thus, if apomicts and sexuals are genetically isolated populations, we expect H_β_ = 2, while if they are genetically a single population, we expect H_β_ = 1^46^. If there are genetic subpopulations along the primary succession, we expect H_β_ ≥ 2 for the six different time windows (maximum H_β_ = 6). We present diversity (D) instead of entropy (H), which is the exponent of the entropy (D = e_H_), and corresponds to the number of markers found.

### Ploidy analysis of mother plants and flow cytometric seed screen

The ploidy level of the sampled mother plants was determined by ploidy analysis using flow cytometry within 48 h after collection of the leaf samples, following the two-step method described by Dolezel and colleagues^47^ with minor modifications. A small piece of a *Bellis perennis* (1.72 pg DNA per nucleus) leaf was added as internal standard to the collected leaf material, which was in 50 µL water. 50 µL of 0.2 M citric acid (Fluka, Buchs, Switzerland), 0.01% Triton X-100 (Sigma-Aldrich, Steinheim, Germany) was added to a total volume of 100 µL, and the leaf material was disrupted by shaking it 2 times for 30 sec at 30 Hz using a mixer-mill (MM300, Retsch, Haan, Germany). After bead-beating 100 µL of 0.1 M citric acid (Fluka, Buchs, Switzerland), 1% Triton X-100 (Sigma-Aldrich, Steinheim, Germany) were added and mixed by inverting the plates to achieve a concentration of 0.1 M citric acid and ca. 0.5% Triton-X-100 in a total volume of 200 µL. The solution was filtered through fritted deep well plates (Nunc, Thermo Scientific, Wohlen, Switzerland) into 96-well V-bottom plates (Sarstedt, Numbrecht, Germany). Nuclei were collected by centrifugation at 150g for 5 min at 20°C (Centrifuge 5810R, Eppendorf, Schönebuch, Switzerland). The supernatant was removed and nuclei were resuspended in 40 µL 0.1 M citric acid, 0.5% Triton X-100. 160 µL of staining solution [0.4 M Na_2_HPO_4_ (Merck, Darmstadt, Germany), 5.5 µg/mL 4’,6-diamidino-2-phenylindole (DAPI; Invitrogen, Eugene, Oregon), and 0.2 µL/mL 2-mercaptoethanol (Sigma-Aldrich, Steinheim, Germany)] were added 2 min prior to analysis by flow cytometer robotics (Quanta SC MPL, Beckman-Coulter, Nyon, Switzerland). The run was stopped at a count of 6000 in the defined sample region or latest after 3:40 min runtime. As the haploid (1C) DNA content of *B. perennis* and *H. pilosella* is the same^48^, the ploidy of samples could be calculated by dividing the median of the *H. pilosella* peak by the median of the *B. perennis* peak and multiplied by 2, to account for diploidy of the *B. perennis* internal standard. We considered individuals with a C_x_ ≥ 5.8 as hexaploid. The protocol and analysis were set up and optimized with tetraploid *H. pilosella* plants which’s ploidy was confirmed by chromosome counts (courtesy of Jan Suda, Department of Botany, Charles University and Institue of Botany, Academy of Sciences, Czech Republic).

The flow cytometric seed screen^49^ followed essentially the same procedure^50^. Single seeds were put into 1.2 mL cluster tubes (Thermo Scientific, Wohlen, Switzerland) containing one 3 mm stainless steel bead (Schieritz & Hauenstein AG, Zwingen, Switzerland). 80 µL of 0.1 M citric acid (Fluka, Buchs, Switzerland), 0.1% Triton X-100 (Sigma-Aldrich, Steinheim, Germany) were added. Seeds were disrupted by shaking them 2 times for 3 min at 30 Hz in a mixer mill. The internal *B. perennis* standard was produced separately from the seeds and used to resuspend the nuclei of the samples. We screened up to 12 seeds per mother plant. This enabled us to detect as low as 8% apomixis per plant. Plants scored as sexual have therefore operationally less than 8% apomixis. In total, we screened 1830 seeds coming from 197 individuals.

### Developmental origin of the seeds

The ratios of the ploidies of (1) endosperm to embryo and (2) embryo to mother plant were used for a linear discriminant analysis (LDA) to assign the developmental origin of the seeds to the four offspring types (n + n, 2n + n, n + 0, 2n + 0, Fig. 1). As training set we used manually annotated data from a different experiment (Sailer et al., *unpublished data*). Datasets were considered to be of sufficient quality if the half peak coefficient of variance HPCV < 5% for the ploidy of the mother, and HPCV < 7% for the ploidy of the embryo. Only datasets of sufficient quality were included in the analysis. We used a higher HPCV value as cutoff in the seed screen because the histograms from seeds from the field are noisier than the histograms from leaves. Furthermore, we excluded all individuals from which we had results of sufficient quality from only one seed. The final dataset contained data from 1231 seeds derived from 153 individual mother plants. As not a single n + 0 type offspring was identified, some 2n + 0 offspring were mis-assigned. Therefore, the n + 0 were removed from the training set and the LDA repeated (Supplementary Fig. 1). The LDA had a wrong assignment rate of 3.4% (Supplementary Fig. 1).

### Statistical analyses

First, we tested the effects of succession, position in the patch (extrinsic factors), and ploidy of the mother plant (intrinsic factor) on the frequency of apomicts, the frequency of the four offspring types, fecundity (number of ovules), and fertility (number of mature seeds/number of ovules) of the mother plant. Second, in a separate analysis, we tested the effect of succession on the ploidy of the mother plant. For all response variables, except ploidy of the mother plant, we used the F-test in ANOVA on generalized linear models (glm), which were first fitted in the order of intrinsic factors, followed by environmental (extrinsic) factors. For testing the effects of succession and patch-position on the ploidy of the mother plant, we used a linear model. By backward elimination of non-significant terms, with keeping variables if they were part of significant interactions, we arrived at the final model. We used the family function “quasibinomial” for over- and underdispersed data with the canonical link function “logit”. Furthermore, the model was weighted by the total number of analyzed individual plants or seeds. In case of interactions, we conservatively tested the corresponding term against the interaction term, instead of against the residual term. All statistical analyses were carried out in R^51^. Graphs were produced using the ggplot2 package^52^ and the grid package^51^.

## Acknowledgements

We thank Nuno Pires and Roger Schmid (University of Zurich) for help with fieldwork and Christian Mavris (University of Zurich) for supplying isochrone data for map construction. We further thank Alain Held and Seraina Camenisch for sorting and counting seeds. We are grateful to Joana Bernardes de Assis, Anja Herrmann, Sofia Magarida Nobre, Philipp Schlüter, and Anja Schmidt (University of Zurich), as well as Andrew Lloyd and Levi Yant (Harvard University) for comments on the manuscript. This work was supported by the University of Zurich and a PSC-Syngenta Fellowship Project of the Zurich-Basel Plant Science Center to J.S. and U.G.

## Author Contributions

U.G. and J.S. conceived and supervised the project, C.S., J.S., and U.G. designed experiments and methodology, C.S. collected and analyzed the data, U.G., J.S., and C.S. wrote the manuscript.

## Data availability

Data generated or analysed during this study are included in the Supplementary Information files.

## References

1 Asker, S. J., Lenn. Apomixis in plants. 1st edn, (CRC Press, 1992).

2 Grossniklaus U. M., JM; Gagliano WB. Molecular and genetic approaches to understanding and engineering apomixis: Arabidopsis as a powerful tool in Advances in Hybrid Rice Technology (ed SS; Siddiq Virmani, EA; Muralidharan, K) Ch. 16, 187–211 (International Rice Research Institute, 1998).

3 Tucker, M. R. et al. Sexual and apomictic reproduction in *Hieracium* subgenus *Pilosella* are closely interrelated developmental pathways. Plant Cell 15, 1524–1537, doi:10.1105/tpc.011742 (2003).

4 Koltunow, A. M. G. et al. Sexual reproduction is the default mode in apomictic *Hieracium* subgenus *Pilosella*, in which two dominant loci function to enable apomixis. Plant. J. 66, 890–902 (2011).

5 Grossniklaus, U., Nogler, G. A. & van Dijk, P. J. How to avoid sex: the genetic control of gametophytic apomixis. Plant Cell 13, 1491–1497, doi:10.2307/3871381 (2001).

6 Tomlinson, J. The advantages of hermaphroditism and parthenogenesis. J. Theor. Biol. 11, 54–58 (1966).

7 Baker, H. G. Self-compatibility and establishment after, long-distance’ dispersal. Evolution 9, 347–349 (1955).

8 Stebbins, G. L. Self fertilization and population variability in the higher plants. Am. Nat. 91, 337–354 (1957).

9 Darlington, C. D. Apomixis: The escape in Evolution of genetic systems Ch. 20, 157–168 (Olvier and Boyd LTD., 1958).

10 Lynch, M. Destabilizing hybridization, general-purpose genotypes and geographic parthenogenesis. Q. Rev. Biol. 59, 257–290 (1984).

11 Hörandl, E. The complex causality of geographical parthenogenesis. New. Phytol. 171, 525–538, doi:10.1111/j.1469-8137.2006.01769.x (2006).

12 Vrijenhoek, R. C. & Parker, E. D. Geographical parthenogenesis: general purpose genotypes and frozen niche variation in Lost Sex: The Evolutionary Biology of Parthenogenesis (eds Isa Schön, Koen Martens, & Peter Dijk) Ch. 6, 99–131 (Springer Netherlands, 2009).

13 Cosendai, A.-C. & Hörandl, E. Cytotype stability, facultative apomixis and geographical parthenogenesis in *Ranunculus kuepferi* (Ranunculaceae). Ann. Bot. 105, 457–470 (2010).

14 Hörandl, E. & Temsch, E. M. Introgression of apomixis into sexual species is inhibited by mentor effects and ploidy barriers in the *Ranunculus auricomus* complex. Ann. Bot. 104, 81–89 (2009).

15 Chapman, H. M., Parh, D. & Oraguzie, N. Genetic structure and colonizing success of a clonal, weedy species, *Pilosella officinarum* (Asteraceae). Heredity 84, 401–409, doi:10.1046/j.1365-2540.2000.00657.x (2000).

16 Morgan-Richards, M., Trewick, S. A., Chapman, H. M. & Krahulcova, A. Interspecific hybridization among *Hieracium* species in New Zealand: evidence from flow cytometry. Heredity 93, 34–42, doi:10.1038/sj.hdy.6800476 (2004).

17 Krahulec, F. & Krahulcova, A. Ploidy levels and reproductive behaviour in invasive *Hieracium pilosella* in Patagonia. NeoBiota 11, 25–31, doi:10.3897/neobiota.11.1349 (2011).

18 Muller, H. J. Some genetic aspects of sex. Am. Nat. 66, 118–138 (1932).

19 Muller, H. J. The relation of recombination to mutational advance. Mutation Research/Fundamental and Molecular Mechanisms of Mutagenesis 1, 2–9 (1964).

20 Apomixis: evolution, mechanisms and perspectives. 1st edn, (Gantner, A R, 2007).

21 Sailer, C., Schmid, B. & Grossniklaus, U. Apomixis allows the transgenerational fixation of phenotypes in hybrid plants. Curr. Biol. 26, 331–337, doi:https://doi.org/10.1016/j.cub.2015.12.045 (2016).

22 Van Dijk, P., de Jong, H., Vijverberg, K. & Biere, A. in Lost Sex: The Evolutionary Biology of Parthenogenesis (eds Isa Schön, Koen Martens, & Peter Dijk) 475–493 (Springer Netherlands, 2009).

23 Koltunow, A. M. & Grossniklaus, U. Apomixis: a developmental perspective. Annu. Rev. Plant. Biol. 54, 547–574 (2003).

24 Bicknell, R. A., Lambie, S. C. & Butler, R. C. Quantification of progeny classes in two facultatively apomictic accessions of Hieracium. Hereditas 138, 11–20 (2003).

25 van der Hulst, R. G. M. et al. Genetic structure of a population sample of apomictic dandelions. Heredity 90, 326–335, doi:10.1038/sj.hdy.6800248 (2003).

26 Houliston, G. J. & Chapman, H. M. Reproductive strategy and population variability in the facultative apomict *Hieracium pilosella* (Asteraceae). Am. J. Bot. 91, 37–44, (2004).

27 Verhoeven, K. J. F. & Preite, V. Epigenetic variation in asexually reproducing organisms. Evolution 68, 644–655 (2014).

28 Greilhuber, J., DoleŽEl, J., LysÁK, M. A. & Bennett, M. D. The origin, evolution and proposed stabilization of the terms ‘genome size’ and ‘C-value’ to describe nuclear DNA contents. Ann. Bot. 95, 255–260 (2005).

29 Catanach, A. S., Erasmuson, S. K., Podivinsky, E., Jordan, B. R. & Bicknell, R. Deletion mapping of genetic regions associated with apomixis in *Hieracium*. P. Natl. Acad. Sci. 103, 18650, doi:10.1073/pnas.0605588103 (2006).

30 Okada, T., Catanach, A. S., Johnson, S. D., Bicknell, R. A. & Koltunow, A. M. An *Hieracium* mutant, loss of apomeiosis 1 (loa1) is defective in the initiation of apomixis. Sex. Plant Reprod. 20, 199–211, doi:10.1007/s00497-007-0057-5 (2007).

31 Rutishauser, A. in Pseudogamie und Polymorphie in der Gattung Potentilla 267–424 (University of Illinois in Urbana-Champaign, 1948).

32 Harlan, J. R. & deWet, J. M. J. On Ö. Winge and a Prayer: The origins of polyploidy. Bot. Rev. 41, 361–390 (1975).

33 Burga, C. A. Vegetation development on the glacier forefield Morteratsch (Switzerland). Appl. Veg. Sci. 2, 17–24 (1999).

34 Burga, C. A. et al. Plant succession and soil development on the foreland of the Morteratsch glacier (Pontresina, Switzerland): Straight forward or chaotic? Flora 205, 561–576 (2010).

35 Mráz, P. Mentor effects in the genus *Hieracium* s.str. (Compositae, Lactuceae). Folia Geobot. 38, 345–350 (2003).

36 Gadella, T. W. J. Cytology and the mode of reproduction of some taxa of *Hieracium* subgenus *Pilosella*. P. K. Ned. Akad. C Biol. 87, 387–399 (1984).

37 Sailer, C., Schmid, B., Stöcklin, J. & Grossniklaus, U. Sexual *Hieracium pilosella* plants are better inter-specific, while apomictic plants are better intra-specific competitors. Perspect. Plant Ecol. 16, 43–51 (2014).

38 Albrecht, M., Riesen, M. & Schmid, B. Plant–pollinator network assembly along the chronosequence of a glacier foreland. Oikos 119, 1610–1624, doi:10.1111/j.1600-0706.2010.18376.x (2010).

39 Bicknell, R. A. & Koltunow, A. M. Understanding apomixis: recent advances and remaining conundrums. Plant Cell 16, S228 (2004).

40 Hörandl, E. et al. Apomixis is not prevalent in subnival to nival plants of the European Alps. Ann. Bot. 108, 381–390 (2011).

41 Winkler, E. & Stöcklin, J. Sexual and vegetative reproduction of *Hieracium* pilosella L. under competition and disturbance: a grid-based simulation model. Ann. Bot. 89, 525–536 (2002).

42 Mráz, P., Šingliarová, B., Urfus, T. & Krahulec, F. Cytogeography of *Pilosella officinarum* (Compositae): altitudinal and longitudinal differences in ploidy level distribution in the Czech Republic and Slovakia and the general pattern in Europe. Ann. Bot. 101, 59–71 (2007).

43 Cryospheric Commission (EKK) of the Swiss Academy of Sciences (SCNAT). The Swiss glaciers. Report No. 1–5, (2012).

44 Okada, T. et al. Chromosomes carrying meiotic avoidance loci in three apomictic eudicot *Hieracium* subgenus *Pilosella*; species share structural features with two monocot apomicts. Plant Physiol. 157, 1327, doi:10.1104/pp.111.181164 (2011).

45 Zini, E. & Komjanc, M. Identification of microsatellite markers in *Hieracium pilosella* L. Conserv. Genet. 9, 487–489 (2008).

46 Jost, L. Entropy and diversity. Oikos 113, 363–375, doi:10.1111/j.2006.0030-1299.14714.x (2006).

47 Doležel, J., Greilhuber, J. & Suda, J. Estimation of nuclear DNA content in plants using flow cytometry. Nat. Prot. 2, 2233–2244, doi:10.1038/nprot.2007.310 (2007).

48 Suda, J. et al. Genome size variation and species relationships in *Hieracium* Sub-genus *Pilosella* (Asteraceae) as inferred by flow cytometry. Ann. Bot. 100, 1323–1335 (2007).

49 Matzk, F., Meister, A. & Schubert, I. An efficient screen for reproductive pathways using mature seeds of monocots and dicots. Plant. J. 21, 97–108 (2000).

50 Sailer, C., Schmidt, A. & Grossniklaus, U. Determination of the developmental origin of seeds containing endosperm using flow cytometric analysis. Bio-protocol 5, e1484, doi:10.21769/BioProtoc.1484 (2015).

51 R: a language and environment for statistical computing (R Foundation for Statistical Computing, Vienna, Austria, 2016).

52 Wickham, H. ggplot2: elegant graphics for data analysis. 1st edn, (Springer-Verlag New York, 2009).

