## Supplementary Fig. 1 for "Dynamics of apomictic and sexual reproduction during primary succession on a glacier forefield in the Swiss Alps"

#### Supplementary Figures

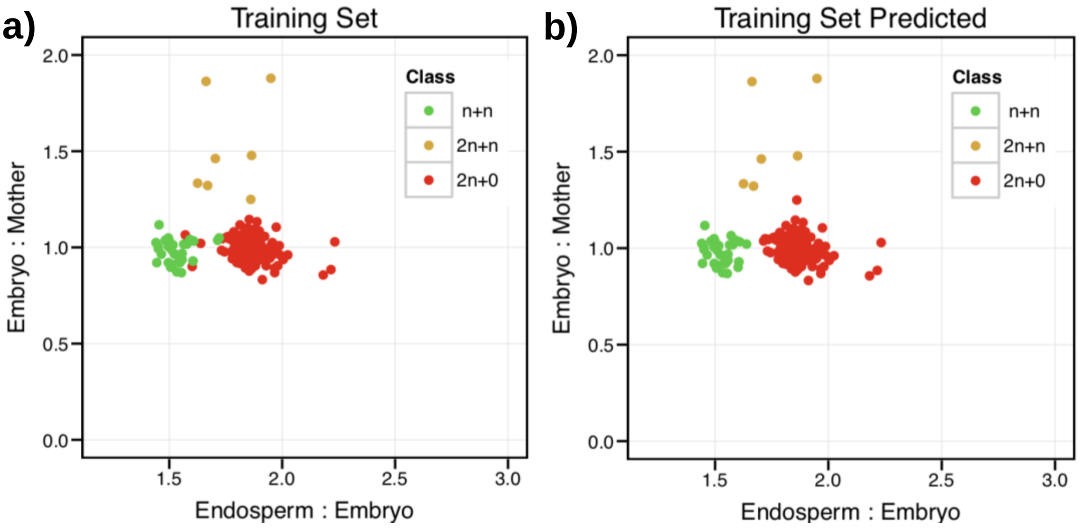


c)

| | n+n | 2n+n | 2n+0 |
|---|---|---|---|
| n+n | 188 | 0 | 12 |
| 2n+n | 0 | 70 | 0 |
| 2n+0 | 4 | 0 | 196 |

**Supplementary Figure 1.** Linear discriminant analysis to distinguish the developmental pathway via which a seed was produced. **a)** Training set used for linear discriminant analysis. The offspring classes were assigned manually based on the flow cytometric histograms. **b)** Linear discriminant analysis of the training set using the identified parameters. **c)** Number of correctly and wrongly assigned offspring.

### Data generated and analysed

**Dataset_Morteratsch** Dataset containing the phenotyping information of all sampled individuals.

**Dataset_GT_LOA267** Genotype matrix of marker LOA267. Column names are binned fragment sizes.

**Dataset_GT_SSR3** Genotype matrix of marker SSR3. Column names are binned fragment sizes.

**Dataset_GT_SSR42** Genotype matrix of marker SSR42. Column names are binned fragment sizes.

**Dataset_GT_SSR87** Genotype matrix of marker SSR87. Column names are binned fragment sizes.
